# In silico λ-dynamics predicts protein binding specificities to modified RNAs

**DOI:** 10.1101/2024.01.26.577511

**Authors:** Murphy Angelo, Wen Zhang, Jonah Z. Vilseck, Scott T. Aoki

## Abstract

RNA modifications shape gene expression through a smorgasbord of chemical changes to canonical RNA bases. Although numbering in the hundreds, only a few RNA modifications are well characterized, in part due to the absence of methods to identify modification sites. Antibodies remain a common tool to identify modified RNA and infer modification sites through straightforward applications. However, specificity issues can result in off-target binding and confound conclusions. This work utilizes in silico λ-dynamics to efficiently estimate binding free energy differences of modification-targeting antibodies between a variety of naturally occurring RNA modifications. Crystal structures of inosine and N6-methyladenosine (m^6^A) targeting antibodies bound to their modified ribonucleosides were determined and served as structural starting points. λ-Dynamics was utilized to predict RNA modifications that permit or inhibit binding to these antibodies. In vitro RNA-antibody binding assays supported the accuracy of these in silico results. High agreement between experimental and computed binding propensities demonstrated that λ-dynamics can serve as a predictive screen for antibody specificity against libraries of RNA modifications. More importantly, this strategy is an innovative way to elucidate how hundreds of known RNA modifications interact with biological molecules without the limitations imposed by in vitro or in vivo methodologies.

## Introduction

Biology has an RNA complexity problem. Cells must make sense of a vast sea of RNAs that function as protein code, regulatory molecules, enzymes, scaffolds, and other biological tools. Furthermore, the 4 canonical RNA bases can be enzymatically modified into new chemical structures that change their ability to base pair, form secondary structure, and interact with RNA-binding proteins (1). These chemical additions can be as small as a single methyl group or as large as a sugar moiety. Over 140 RNA modifications have been identified across all three kingdoms of life (1). RNA modifications are prevalent in biology and function as an epigenetic code to regulate development (2), respond to infectious diseases (3), and are involved in cancer progression (4). Their combinatorial complexity highlights how individual or collections of RNA modifications may alter an RNA’s fate or function. A current challenge is the development of methods to identify all modification sites to decipher the roles of these RNA modifications in biology.

A variety of methods can identify a few RNA modification sites. For example, chemical treatment can identify m^6^A (e.g. GLORI (5)) and pseudouridine (e.g. pseudo-seq (6)) by taking advantage of chemistries that affect a modified base differently than an unmodified base. Direct RNA nanopore sequencing can also identify specific modifications like m^6^A (7–17) through differences in electrical current perturbations as the modified RNA transverses the sequencing pore. Both strategies, however, require tailor-made approaches to accommodate each RNA modification’s unique biochemical characteristics. Furthermore, without employing enrichment strategies, low abundance modifications remain difficult to detect. Adaptable methods are needed to elucidate the full breadth of modified RNAs found in living organisms.

A common, versatile identification strategy uses antibodies to immunoprecipitate modified RNAs (18). These enriched RNAs are then sequenced to identify RNA targets and infer modification sites. Immunoprecipitation and sequencing methods are well established with straightforward workflows, and enrichment permits identification of less prevalent modification sites. Indeed, much of the work determining the modification sites of N6-methyladenosine (m^6^A, e.g. (19,20)), N1-methyladenosine (m^1^A, e.g. (21–24)), 5-methylcytosine (m^5^C, e.g. (25,26)), and others have relied on antibodies.

Antibodies can become de novo RNA-binding proteins through adaptive immunity. Immunoglobulin G (IgG) antibodies are comprised of two heavy and two light polypeptide chains that assemble a pair of six hypervariable complementary-determining region (CDR) loops at their antigen recognition interface (27–29). Antibodies recognize a variety of antigens through CDRs that vary in amino acid length and composition. How antibodies recognize proteins is well studied (30), but how antibodies recognize modified RNAs is less clear. A polyinosine-antibody crystal structure was determined bound to various nucleotides (31). Closer inspection of the structure reveals a large, suitably configured pocket adjacent to the bound nucleotide (**Fig S1**), suggesting that the antibody may have specificity toward nucleic acid, not single bases. Regardless, the lack of antibody structures targeting other modified bases limits insights into how antibodies recognize RNA modifications.

The success of using antibodies for RNA modification site identification depends on the quality of the antibody (32,33). Antibodies with low specificity have assigned erroneous biochemical functions to RNA modifications. For example, published studies reached differing conclusions regarding the mechanism of the m^1^A modification. Two studies found m^1^A prevalent in the 5’ ends of mRNA (23,24), suggesting that the modification enhances translation (24), while contrasting studies reported it as rare in mRNA (21,22). In the former studies, it was later discovered that the antibody used for m^1^A RNA enrichment also had affinity towards 7-methylguanosine (m^7^G, (21)), an abundant mRNA 5’ cap modification crucial for cap-dependent translation (34). These false positive site identifications led to incorrect conclusions regarding m^1^A function. Because the identification of RNA targets and their specific modification sites gives insight into their biological and biochemical mechanisms, the development of antibodies with high affinity and high specificity is a key to successfully discovering the biological roles of the many RNA modifications. However, given the large number of RNA modifications and the subtle chemical differences between them, off-targets of RNA modification antibodies will be a continuous, inevitable problem. The current state of RNA chemistry prevents in vitro testing of all known RNA modifications, and thus new methods are required to predict the specificity of RNA modification-targeting antibodies.

Computational approaches have the potential to screen antibodies for their predicted ability to bind modified RNA bases. Physics-based, alchemical free energy calculations are an accurate, rigorous, and cost-effective means to quantify chemical probe interactions with protein structures in silico (35–37). These calculations compute relative binding free energies (ΔΔ*G*_bind_) between two or more molecules by transforming between alternate chemical groups in silico. Because they are at the heart of molecular dynamics simulations, these calculations also provide dynamic structural characterization of macromolecular complexes. With these methods, changes in RNA-protein binding affinities can be monitored as a function of the chemical differences between modified or unmodified RNAs. Hence, modeling different RNA modifications can predict binding selectivity.

λ-Dynamics is an efficient alchemical free energy method that can accurately and rapidly screen hundreds of modified RNAs bound to a protein host. This method holds a key advantage over other in silico strategies in that it can model multiple chemical variations simultaneously within a single simulation (38,39), making it more efficient and higher throughput. In a λ-dynamics calculation, a variable λ parameter allows chemical groups to dynamically scale between “on” and “off” states during a molecular dynamics simulation. Akin to selection in an in vitro competitive binding assay, this dynamic behavior effectively differentiates the varying affinities of target molecules, providing insights into their binding characteristics. Thus, λ-dynamics can rapidly select for the best binders from a library of chemical modifications (40,41). To date, λ-dynamics has accurately measured the relative binding free energy differences of large chemical inhibitor libraries targeting the HIV reverse transcriptase (42–44) and β-secretase 1 (45,46), of mutations at various protein-protein interfaces (47,48), as well as of the folding free energies of mutant T4 lysozyme proteins (49). Notably, chemical probe binding studies with λ-dynamics demonstrated 8- to 30-fold efficiency gains over other conventional free energy calculations (42,45). This equates to months of computational time savings.

The following investigation tested whether λ-dynamics could accurately predict how RNA modifications affected RNA-protein interactions. This work determined the structures of two modified RNA-targeting antibodies bound to inosine and m^6^A, revealing that these antibodies recognize their target ligands similar to other modified RNA binding proteins. The structural models permitted the use of λ-dynamics to perform a computational screen of RNA base modifications bound to inosine and m^6^A antibodies to predict their binding specificities. These in silico binding predictions were verified with in vitro binding assays. Collectively, the results demonstrate how structural biology can be combined with λ-dynamics to predict modified RNA-protein interactions without the limitations imposed by biochemical experiment methodologies.

## Results

The goal was to test whether λ-dynamics could be used as an in silico strategy to accurately probe modified RNA-protein interactions. Antibodies can serve as modified RNA-binding proteins. They are commonly used as reagents to enrich for modified RNAs and determine modification sites in biology (18). Currently, RNA modification targeting antibodies are relatively few in number, have modest affinity toward their targets (32,33), and can have specificity issues that confound biological conclusions (21). An antibody specificity screening method for known RNA modifications will enable a comprehensive view of the RNAs enriched and provide insight into how to improve antibody design.

High-resolution structures of antibodies targeting single modified RNA bases have not been published. An inosine-targeting antibody structure is available (31), but an open pocket adjacent to the nucleoside binding site potentiates the chance of the antibody binding to a dinucleotide substrate (**Fig S1**). To avoid this confounder, additional antibody structures bound to modified ribonucleosides were pursued. The protein sequences of available antibodies were predicted by mass spectrometry and sequencing (see Methods). Recombinant antibodies were produced in cell culture and used to generate antibody fragments (Fabs). Fabs were screened in crystallizing conditions, and crystals were soaked or grown with target nucleoside ligands (see Methods). These efforts lead to the determination of three modified RNA-targeting antibody crystal structures (**Table S1**): one targeting inosine at 1.94 Å and two targeting m^6^A at 2.02 Å and 3.06 Å.

IgG antibodies are composed of heavy and light protein chains, forming 6 variable loops on each arm, or antibody binding fragment (Fab), that typically dictate binding affinity to its target substrate (27–29). In the 1.94 Å inosine and 3.06 Å m^6^A antibody structures, a large, discontinous density was observed at these variable loop regions where a modified purine target nucleoside could be adequately modeled (**Fig 1A,B**). Rather than binding to loops on the periphery, the modified nucleosides bound to a central cavity created by the 6 variable loops between the heavy and light chains (**Fig 1A,B**). Binding of small molecules at this location has been observed in other antibody structures (50). In the third 2.02 Å m^6^A targeting antibody structure, density in this binding pocket was not observed (**Fig S2**). Thus, two structures yielded high-resolution models of how purine modified bases bind to antibodies.

**Fig 1.**
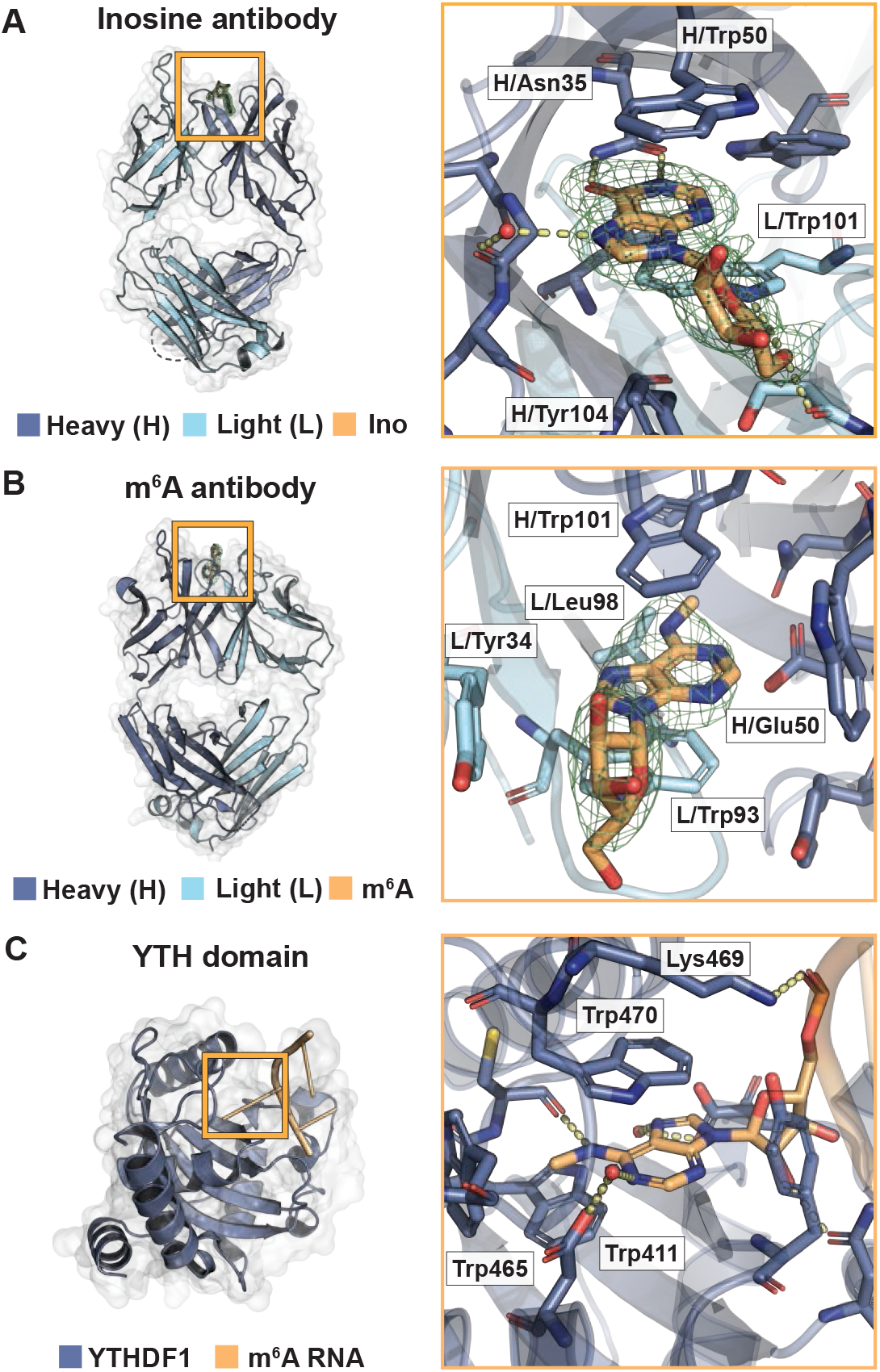
Binding of inosine and m^6^A targeting antibodies mimics RNA-binding proteins. (**A**) Crystal structure of the inosine targeting antibody fragment to 1.94 Å (PDB ID: 8SIP). Overview (left) and magnified (right) rendition of the antibody bound to inosine nucleoside. 1F_o_F_c_ density without ligand in green mesh. Heavy chain (H) in dark blue, light chain (L) in light blue, and inosine in orange. Interacting amino acids include heavy chain residues Asn35, Trp40, Trp50, Gly99, Tyr104, and Leu106 and light chain residues Ser97 and Trp101. Those discussed in the main text are labeled. (**B**) Crystal structure of the m^6^A-targeting antibody fragment to 3.06 Å (PDB ID: 8VEV). Labeling same as in (A), except m^6^A nucleoside in orange. Interacting amino acids include heavy chain residues Trp33, Asn35, Glu50, Tyr61, Trp101, and Phe105 and light chain residues Tyr34, Trp93, and Leu98. Those discussed in the main text are labeled. (**C**) Structure of a YTH bound to m^6^A (YTHF1, PDB ID: 4RCJ). Residues in dark blue. m^6^A in orange. Interacting amino acids include Tyr397, Asp401, Trp411, Cys412, Asn441, Trp465, Lys469, Trp470, and Asp507. Those discussed in the main text are labeled.

Small molecule antibodies are selected through adaptive immunity to target a particular hapten (51). Thus, antibodies become RNA-binding proteins through adaptation and can inform on how biology designs a protein to bind an RNA modification de novo. Modified RNA-binding proteins provide exemplary examples of potential binding architecture. For example, the YTH domains bind to m^6^A with high specificity (52). This domain arranges its side chains to 1) create a specificity pocket for the parent base and modification, 2) bind the nucleobase through π-π stacking, and 3) line the pocket periphery with positively charged side chains to accommodate the negatively charged RNA phosphate backbone (**Fig 1C**). Antibodies targeting modified RNAs might also mimic this strategy. Alternatively, they might use a collection of novel binding strategies, each selected randomly through adaptive immune selection.

The inosine and m^6^A antibody structures both bound to their modified ribonucleoside ligands similarly to other RNA-binding proteins. To specify the modified base, the inosine targeting antibody used an asparagine to select for the O6 oxygen and N1 nitrogen of the inosine nucleobase (**Fig 1A**). The m^6^A-targeting antibody created a hydrophobic pocket to accommodate the methyl group (**Fig 1B**) and a glutamate side chain to hydrogen bond with the adenosine nucleobase N1 nitrogen (**Fig 1B**). Interestingly, glutamate side chain coordination is also observed in some YTH domains that bind m^6^A (**Fig 1C**, (53)). Both antibodies used paired tryptophans to create a slot for favorable π-π stacking and a tyrosine for ribose ring interactions (**Fig 1A,B**). However, these tryptophans and tyrosine came from differing variable loops in each antibody and are organized differently in their central antibody binding pocket (**Fig 1A,B**). The difference in binding pocket organization potentially reflects how these two antibodies were isolated from different animals with separate adaptive immune responses. In sum, the antibody-ligand structures revealed that these two antibodies use similar strategies to bind their modified base targets that may permit differentiation between unmodified base counterparts.

The quality of the structures enabled predicting in silico how these antibodies may interact with other RNA nucleobases. There are over 140 different RNA modifications identified in biology, many of which are not available as commercial reagents or lack protocols to synthesize in vitro. A library of 44 modified and 4 unmodified nucleobases was selected based on published thermodynamic parameters for RNA modifications in the CHARMM force field (54) and their commercial availability for experimental testing in vitro (**Fig S3**). λ-Dynamics was used to assess differences in relative binding free energies between inosine or m^6^A versus each library nucleobase when bound to their respective antibodies (see Methods, **Fig 2**, and **Fig S4**). During the simulations, some of the modified nucleosides unbound from the antibody (**Fig S5**), presumably due to having poor binding affinity or steric clashes, and were removed from further study (**Table S2** and **S3**). Similar to previously performed studies (42–44,47–49), relative binding free energies (ΔΔ*G*_bind_) were calculated for the nucleosides that remained antibody bound. Examples of the results obtained are shown (**Fig 3** and **4**) with full results reported in the Supplement (**Table S2 and S3**). A positive ΔΔ*G*_bind_ value indicates poorer binding and a negative value suggests enhanced binding when compared to the native inosine or m^6^A base. As a control, inosine and m^6^A modified bases were perturbed into an identical but distinct copy of themselves within their respective antibody complexes. These free energy differences were near zero (**Fig 3A and 4**), as expected of a base replacing itself, and indicated that the λ-dynamics calculations were working correctly.

**Fig 2.**
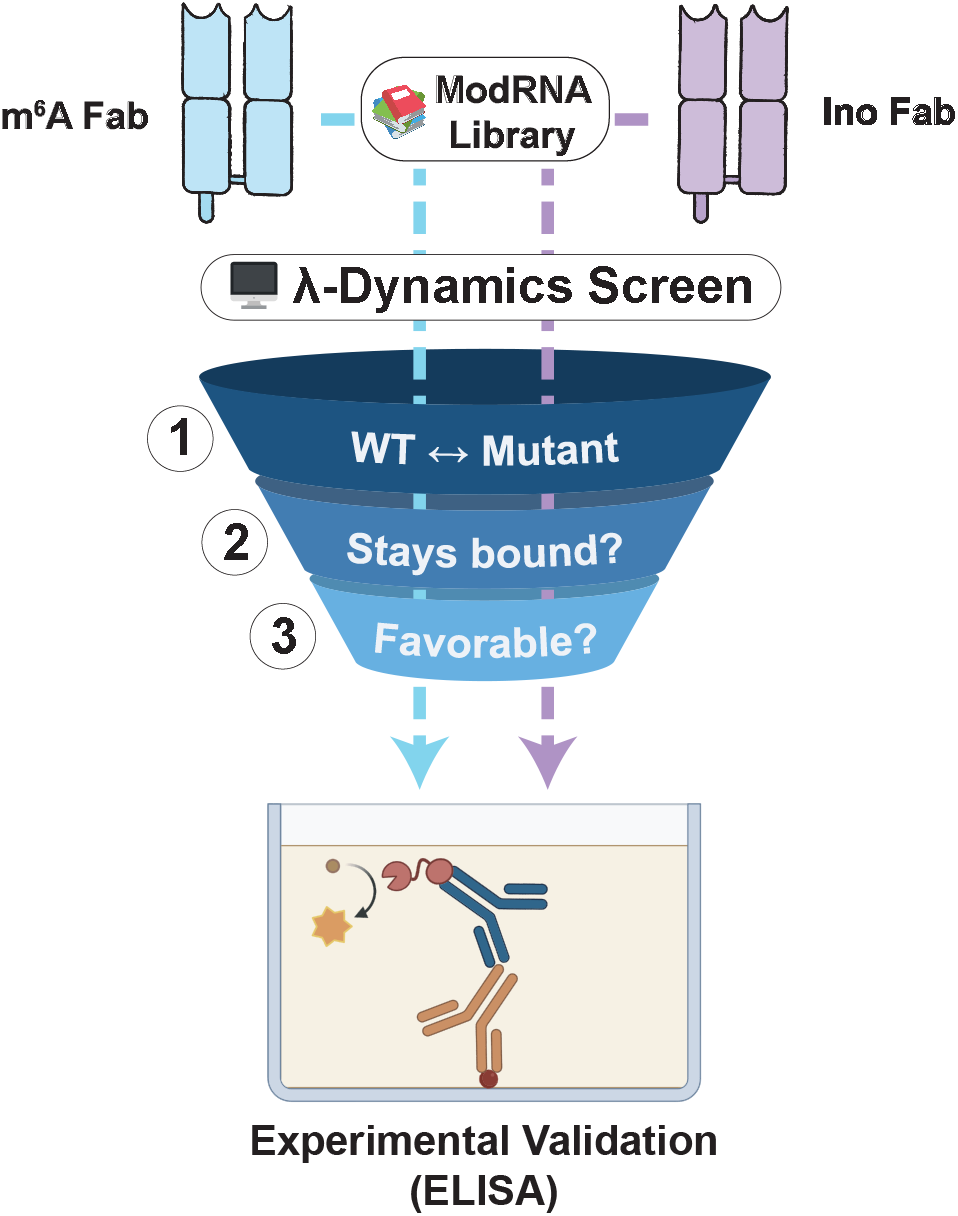
In silico λ-dynamics workflow for screening potential binders to the inosine and m^6^A antibodies. A three-step process was used to filter candidates from a library of 48 ribonucleosides for in vitro antibody binding validation. (1) For each mutant library candidate, a λ-dynamics simulation was conducted to calculate a relative binding free energy between the mutant and its respective native ribonucleoside ligand (inosine or m^6^A). (2) All ribonucleosides that unbound during these simulations were deemed unfavorable and excluded from further processing. (3) Mutant bases with relative binding free energies deemed favorable (ΔΔ*G*_bind_ ≤ −0.7 kcal/mol) were selected for *in vitro* validation with binding assays based on commercial availability.

**Fig 3.**
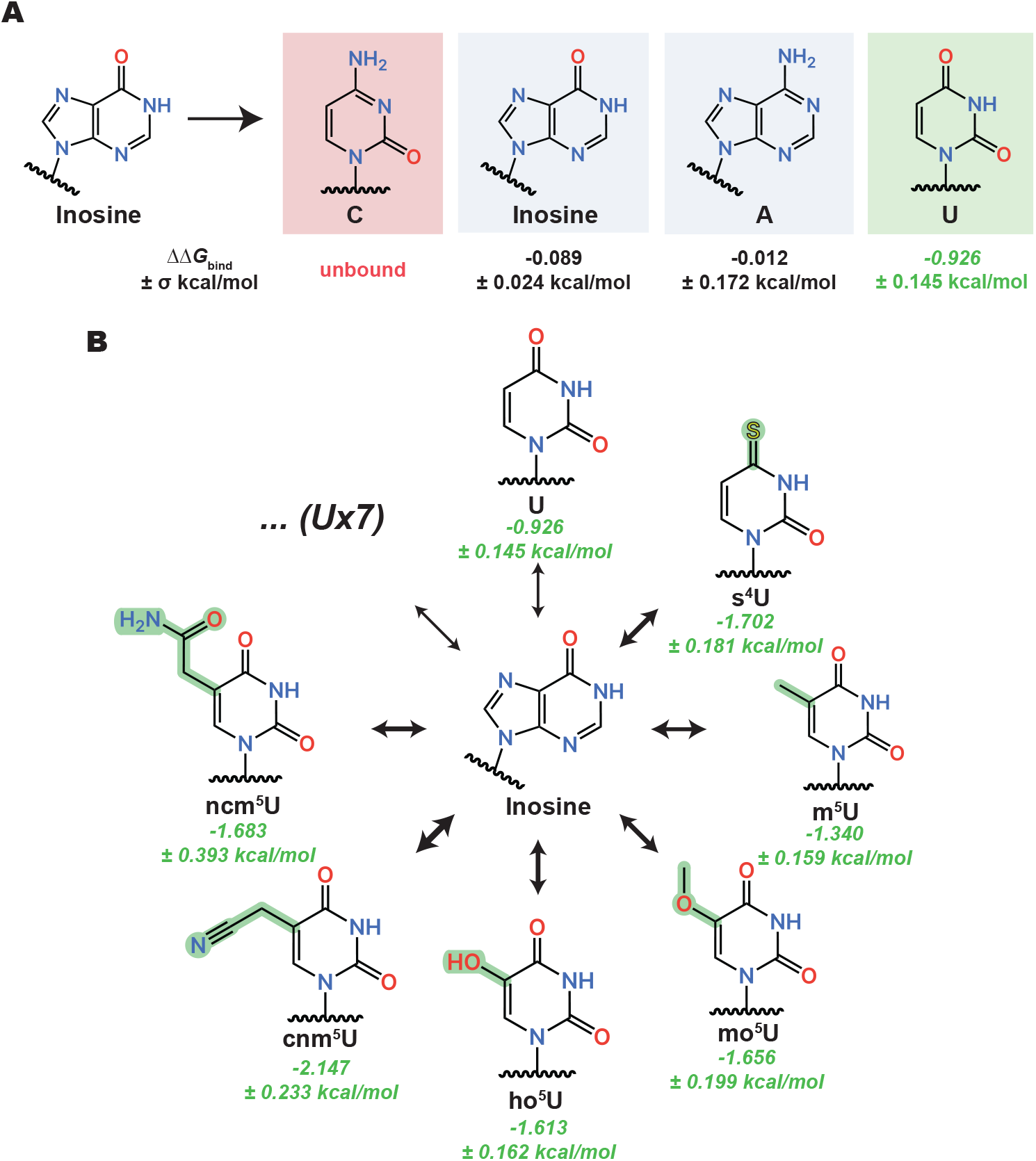
Highlighted binding trends from the inosine antibody λ-dynamics screen. (**A**) λ-Dynamics predicts loss of binding (red) for cytidine (C), no change in binding (grey) for inosine and adenosine (A), and enhancement of binding (green) for uridine (U). Estimated relative binding free energies (ΔΔ*G*_bind_) and uncertainties (±*σ*) are listed. (**B**) The predicted inosine antibody promiscuity for U generalizes to many of its derivatives. Estimated relative binding free energies and uncertainties are listed in green. The thickness of each equilibrium arrow is proportional to the favorability of the corresponding transition. Seven other uridine derivatives (Ux7) showed enhanced binding but are not depicted. See **Table S2** for a complete list.

**Fig 4.**
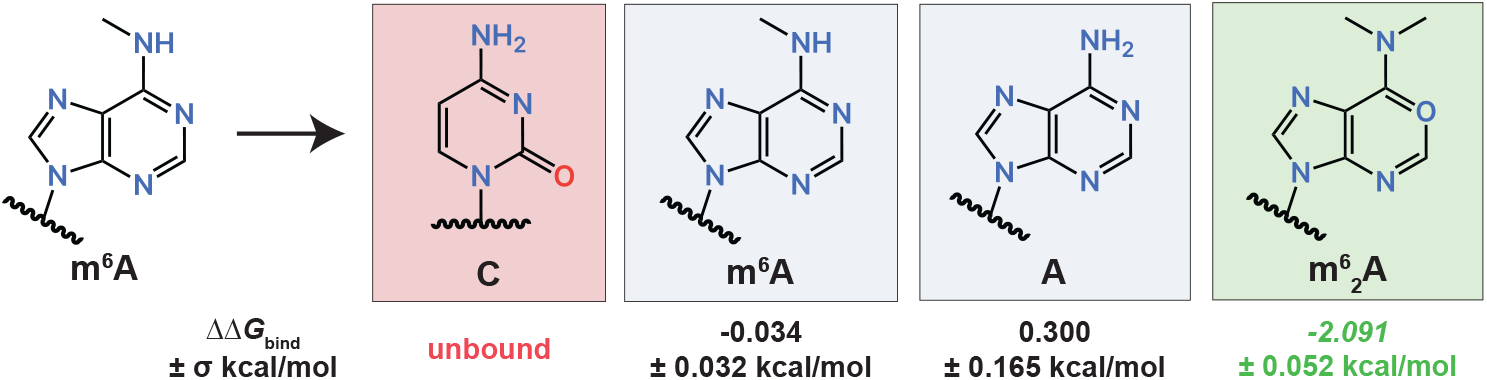
Highlighted binding trends from the m^6^A antibody λ-dynamics screen. λ-Dynamics predicts loss of binding (red) for cytidine (C), no change in binding (grey) for m^6^A and adenosine (A), and enhancement of binding (green) for m^6^_2_A. Estimated relative binding free energies (ΔΔ*G*_bind_) and uncertainties (±*σ*) are listed. See **Table S3** for a complete list.

λ-Dynamics predicted differing specificities and off-targets for these two antibodies. The inosine antibody had many predicted off-targets that included uridine (**Fig 3A**) and uridine modifications (**Fig 3B**). Inspection of the models revealed that hydrogen bonding of the asparagine side chain to the O6 oxygen in inosine could be satisfied by the O4 oxygen in uridine (**Fig S6A**). Many uridine modifications had an O4 oxygen available for hydrogen bonding, potentially explaining why related molecules all had higher predicted binding affinities in the λ-dynamics calculations. In contrast, cytidine and adenosine were not predicted to enhance binding (**Fig 3A** and **Table S2**). Both nucleosides have nitrogens at similar positions, potentially making the pocket less favorable for these bases to interact by removing hydrogen bonding. Finally, a further inspection of the structures revealed a larger binding pocket in the inosine versus the m^6^A antibody binding pocket (**Fig 1A,B**). This larger pocket may accommodate a greater variety of shapes and sizes, increasing the propensity for off-targets. Thus, λ-dynamics predicted the inosine antibody to have many off-targets in this modestly sized ribonucleoside library.

In contrast to the inosine antibody, λ-dynamics predicted that the m^6^A antibody had relatively few off-targets (**Table S3**). As discussed previously, the binding pocket was smaller (**Fig 1A,B**) and required a N1 nitrogen on the nucleobase for hydrogen bonding (**Fig 1B**). Along with m^6^A, a few adenosine bases were predicted to bind (**Fig 4** and **Table S3**), including adenosine and N6,N6-dimethyladenosine (m^6^_2_A), a dimethyl modification at the N6 nitrogen position (**Fig S6B,C**). Closer inspection of the structure revealed that the hydrophobic pocket had enough space to accommodate a second methyl group (**Fig S6C**). Similar to the inosine antibody, cytidine was predicted to be a poor binder with a high, positive free energy difference (**Fig 4**). In summary, the m^6^A antibody had fewer off-targets compared to the inosine antibody but still was predicted to bind to nucleosides other than m^6^A.

While λ-dynamics has demonstrated accuracy with modeling protein-protein and protein-small molecule binding interactions (42–48), it has so far been untested with respect to reproducing protein-RNA interactions. To evaluate our in silico predictions in vitro, Enzyme-Linked Immunosorbent Assays (ELISAs) were used to probe the binding of inosine and m^6^A antibodies to target and off-target RNA bases. RNAs were synthesized through solid-state chemistry (see Methods) to create biotin-labeled oligomers of inosine, adenosine, uridine, and cytidine to test the inosine antibody binding. Cytidine oligos with single base changes of adenosine, m^6^A, and m^6^_2_A were synthesized to test the m^6^A antibody binding. The biotin-labeled oligos were bound to wells coated with a streptavidin derivative. Wells without oligo served as a background control. After oligo incubation and washing, the inosine and m^6^A antibodies were incubated at varying concentrations. Bound inosine and m^6^A antibodies were detected with a secondary horseradish peroxidase (HRP) conjugated antibody that targeted mouse IgG. No inosine or m^6^A antibody wells were used to control for secondary antibody background. The presence of secondary antibody was detected with an HRP chromogenic substrate, with the absorbance measured as an indirect reading for inosine or m^6^A antibody binding.

The inosine and m^6^A antibody in vitro binding results agreed with the λ-dynamics predictions (**Fig 5**). The inosine antibody bound to inosine and uridine oligos (**Fig 5A**), although inosine binding was observed at much lower antibody concentrations. In contrast, the inosine antibody did not bind to adenosine or cytosine oligos (**Fig 5A**). Likewise, the m^6^A antibody bound to m^6^A containing cytidine oligos but bound poorly to cytidine only (**Fig 5B**), as expected. As λ-dynamics predicted, the m^6^A antibody bound to an m^6^_2_A-containing oligo (**Fig 5B**). The antibody also bound to an adenosine-containing oligo (**Fig 5B**) but to a lesser degree than m^6^A. Regardless, the in vitro binding results matched the predictions of λ-dynamics, supporting the accuracy of this in silico method to identify modified RNA-protein interactions.

**Fig 5.**
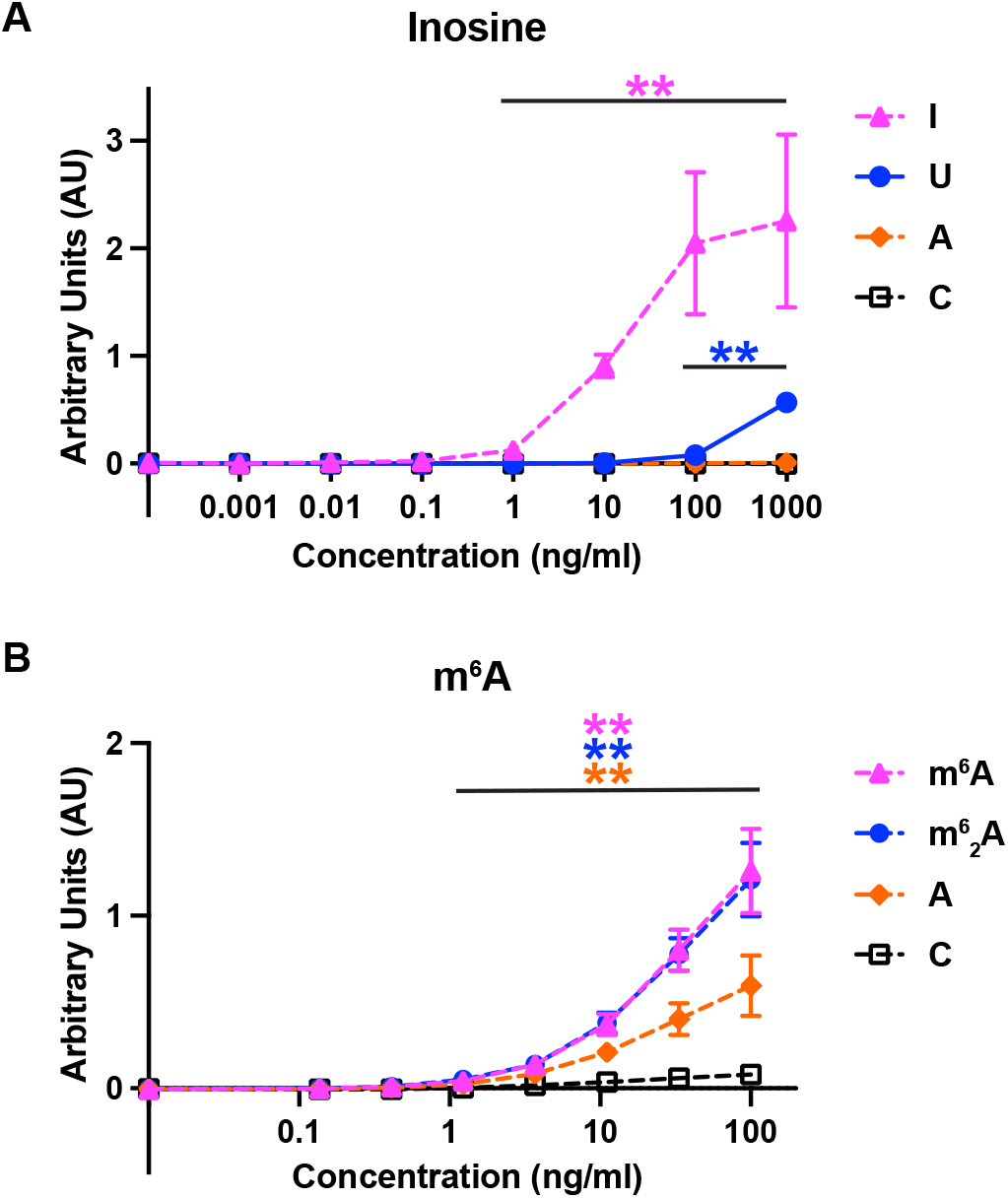
ELISA binding assay results confirmed λ-dynamics predictions of antibody off-targets. (**A**) Absorbance units reported by ELISA indicating the binding affinity of inosine antibody to inosine (I), uridine (U), adenosine (A), and cytidine (C) over varying protein concentrations. Double asterisks (**) denote a p-value < 0.01. Inosine serves as a positive control. In line with λ-dynamics predictions, U identified as an off-target while A and C demonstrated negligible binding. (**B**) Absorbance units reported by ELISA indicating the binding affinity of m^6^A antibody to m^6^A, m^6^_2_A, adenosine (A), and cytidine (C) at varying protein concentrations. Double asterisks (**) denote a p-value < 0.01. m^6^A serves as a positive control. Again, matching λ-dynamics predictions, m^6^_2_A and A are identified as off-targets while C demonstrated negligible binding. All p-values calculated are available in **Fig S6D,E**.

## Discussion

With hundreds of RNA modifications identified in biology, new methods are required to determine the sites of each of these chemical changes to determine their functions. Antibodies targeting RNA modifications are a versatile tool to enrich and determine modification sites, but their reliability hinges upon their accuracy. To this end, inosine and m^6^A antibody structures bound to their modified ribonucleoside targets were determined to high resolution. These structures then facilitated the use of λ-dynamics, an in silico free energy calculation, to estimate how the antibodies may bind other unmodified and modified RNA bases, with worsened, neutral, or enhanced binding affinities. λ-Dynamics predictions matched well with in vitro binding assay results, supporting the accuracy of using this computational approach to measure untested RNA-protein interactions. In its simplest application and as performed in this work, the method can be used to determine off-target RNA base interactions with antibodies used for modified RNA enrichment and site identification. But the strategy holds greater promise to inject insight into the biochemical mechanisms of RNA modifications by determining how any modified RNA, commercially available for biochemical investigation or not, may interact with proteins and other molecules (**Fig 6**).

**Fig 6.**
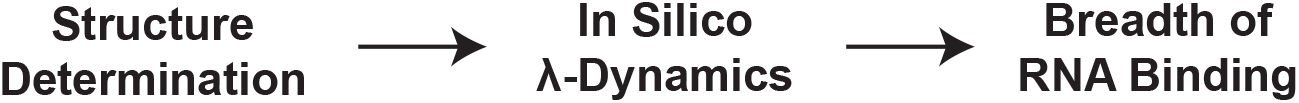
Proposed strategy to predict how proteins bind canonical and modified RNA. (1) Starting with an RNA-protein structural model, (2) an in silico λ-dynamics screen can be conducted to assess the favorability of the protein’s interactions with a complete range of RNA bases. (3) This approach provides an economical and effective means to explore the full extent of a protein’s RNA-binding capabilities that can be tested further in vitro.

The determined antibody structures targeting modified purines revealed identical binding strategies toward their respective modified RNA bases, reminiscent of modified RNA-binding proteins. Each antibody had a specificity pocket and used tryptophans to create a slot for π-π stacking with the nucleobase. Only one of these tryptophans had a similar sequence position between the two antibodies. The other came from a separate loop, leading to RNA binding in completely different orientations. These antibodies were created through adaptive immunity, supporting the notion that mimicking modified base RNA-binding proteins by creating a specificity pocket and using π-π stacking for nucleobase interactions is a competent way to bind a modified nucleobase. Thus, convergent adaption may have led both purine-targeting antibodies to follow a similar binding strategy as modified RNA-binding proteins. The results lead to the speculation that all modified RNA-targeting antibodies bind to their targets similarly. Examples of pyrimidine-targeting antibody structures will be necessary to further probe this concept.

Antibodies are heavily used reagents to enrich modified RNA for sequencing and site identification. This strategy has been used to identify sites of many different RNA modifications to deduce their biological and biochemical mechanisms. Regardless of new methodologies to determine RNA modification sites, antibodies will continue to be used to enrich for less abundant modifications. Thus, antibody binding to off-target RNA modifications will continue to be a problem in research. The chemical similarities between many RNA modifications make antibody specificity an expected complication. This work demonstrates how λ-dynamics is a viable in silico tool to determine potential RNA off-targets of antibodies. The method does not require the availability of modified nucleosides, RNA oligomers, or other in vitro reagents that are currently unavailable. With an accurate, high-resolution structural model, λ-dynamics can test the full breadth of RNA modifications in biology. Additionally, λ-dynamics has previously investigated the effects of protein mutations on binding (47,48). The method can thus be used to rationally design antibodies for improved binding specificity and affinity.

This is the first study to use λ-dynamics to probe nucleic acid-protein interactions via nucleic base perturbations. Other in silico molecular modeling and free energy methods have been employed to study nucleic acid-protein interactions, including predictions of DNA binding to proteins (55) and probing mutations in DNA-protein complexes (56,57). λ-Dynamics has several key attributes that make it advantageous over other in silico calculations. First, λ-dynamics enables multiple modified bases to be calculated within a single simulation. This can drastically improve efficiency over other free energy methods that can only investigate a single perturbation at a time, therefore requiring many simulations to study multiple perturbations. Second, λ-dynamics can simultaneously sample modifications at multiple sites within a chemical system. This enables base changes at different RNA sequence positions to yield free energy results for multiple modification combinations. There are limitations to λ-dynamics as well. Many of the calculated free energy differences, such as with uridine bound to the inosine antibody (**Fig 3A**) or with m^6^_2_A bound to the m^6^A antibody (**Fig 4**), predicted greater enhancement of binding than what was observed in vitro (**Fig 5**). The starting models for the λ-dynamics calculations were based on the crystal structures of antibody fragments bound to nucleosides, but binding was tested in vitro with RNA oligos. This omission of the RNA phosphate backbone from the model, as well as the potential for sporadic self-associations or secondary structures in the unbound oligo, may have impacted the true binding values. Additional work probing RNA-protein interactions with λ-dynamics will undoubtedly improve the simulations. Moreover, the refinement of molecular dynamics force fields, particularly with respect to nucleic acids, is a bustling area of research, and future advancements promise to further enhance the accuracy of these classical simulations.

While hundreds of RNA modifications have been identified, only a few dozen are available for experimental testing in vitro. Novel methods must be developed to examine how all modifications affect molecular interactions to decipher their biological mechanisms. This study establishes a workflow for using λ-dynamics to probe nucleic acid-protein interactions in silico (**Fig 6**). The combinatorial efficiency of λ-dynamics enables rapid in silico examination of currently known and newly discovered RNA modifications. With high-resolution structures of nucleic acid-protein complexes, modified and unmodified nucleoside bases can be probed to explore how chemical changes to RNA affect protein binding interactions. This computational approach can be used for DNA or RNA and is not limited by available chemistry. The work presented demonstrates how this strategy can probe for the specificity of antibodies. Future work can utilize this method to test how hundreds of RNA modifications affect their molecular interactions with any RNA-binding protein or other nucleic acids, delivering novel insights into their molecular functions.

## Materials and Methods

### Recombinant antibodies

Commercial antibodies targeting inosine and m^6^A were sequenced by Abterra Biosciences (San Diego, CA) (58–60). Briefly, the antibodies were fragmented and submitted for MS/MS mass spectrometry. The data was then analyzed to predict the probable antibody sequence. Full-length monoclonal antibodies (mAb) and antibody fragments (Fab) were produced recombinantly in human cells by Sino Biological (Wayne, PA). Fabs were made from mAbs by papain protease digestion, Fc removal by protein A, and size exclusion chromatography. All mAbs and Fabs were shipped and stored in phosphate buffered saline (PBS; 137 mM NaCl, 2.7 mM KCl, 10 mM Na_2_HPO_4_, 1.8 mM KH_2_PO_4_).

### Crystallography

Recombinant Fabs were concentrated to approximately 3-5 mg/ml and sitting drop crystal trays were set with an Oryx4 (Douglas Instruments; Hungerford, United Kingdom). The m^6^A Fab was set up without and with 1 mM m^6^A nucleoside (MedChemExpress, HY-N0086). Crystals were observed by 4 weeks in the following conditions: 1) the inosine Fab in 50 mM Tris pH 8.3, 15% PEG 4000, 0.1 mM EDTA; 2) the m^6^A Fab only in 20% (v/v) PEG 2K, 0.2 M MgCl2, 100 mM Tris pH 8.0; and 3) m^6^A Fab with 1 mM m^6^A nucleoside in 0.17 M ammonium sulfate, 25.5% (w/v) PEG 4000. The inosine and m^6^A Fab only crystals were incubated in freezing conditions (inosine: 21% PEG 4K, 50 mM Tris pH 8.3, 0.1 mM EDTA, 20% glycerol, 0.2 mM inosine nucleoside (Sigma, I4125-1G); m^6^A: 20% (v/v) PEG 2K, 0.2 M MgCl_2_, 100 mM Tris pH 8.0, 5-15% (v/v) glycerol, 1 mM m^6^A nucleoside) with addition of 10 mM inosine and 10 mM m^6^A nucleoside for 30-60 minutes prior to freezing, respectively. X-ray diffraction data was collected at Lilly Research Laboratories Collaborative Access Team (LRL-CAT; Argonne National Laboratory; Argonne, IL) and ESRF ID30B (Life Sciences Collaborative Access Team (LS-CAT) operating at the European Synchrotron Radiation Facility (ESRF); Grenoble, France). Data was collected and processed by Lilly, UW-Madison Crystallography Core, and the authors. All data was indexed, merged, and scaled in XDS/Aimless (61). Space groups were determined in XDS/pointless (61). Model building and refinement were performed in Coot (62) and Phenix (63), respectively. In some of the inosine and m^6^A Fab density maps, a large density was observed at the Fab antigen binding site. The respective modified RNA nucleosides used in crystallization and in freezing modeled well into these densities (**Fig 1A,B**). The final structures and merged reflection files are deposited at wwPDB (wwpdb.org; PDB IDs: 8SIP, 8TCA, 8VEV). Unmerged reflection data were deposited at Integrated Resource for Reproducibility in Macromolecular Crystallography (proteindiffraction.org).

### System setup for molecular modeling

Coordinates for the inosine and m^6^A Fabs were obtained from our Protein Data Bank (PDB) entries 8SIP and 8VEV. Residue flips for His, Glu, and Asn were assessed using the MolProbity webserver (64). Protonation states of titratable residues were assigned based on their predicted pKa values at pH 7.0 using PROPKA (65,66). The protein-nucleoside complexes were then solvated using the CHARMM-GUI Solution Builder (67), requiring a minimum of 10 Å of solvent padding from each face. The resulting cubic water box dimensions were 101 Å per edge for the inosine system and 98 Å per edge for the m^6^A system. Sufficient K^+^ or Cl^−^ ions were added to neutralize the net charge of each system. Additional K^+^, Mg^2+^, and Cl^−^ ions were then added to achieve a final ionic strength of 150 mM KCl and 0–5 mM MgCl_2_. This process was repeated to solvate the individual nucleosides without their respective Fabs, yielding unbound model systems with cubic box dimensions of 30 Å per edge for inosine and 32 Å per edge for m^6^A.

All simulations were performed using the CHARMM molecular simulation package ((68,69), developmental version c47a2) with the Basic λ-Dynamics Engine (BLaDE) on graphics processing units (GPUs) (70). Prior to running molecular dynamics, each system was subjected to 250 steps of steepest descent minimization. Molecular dynamics (MD) simulations were then run in the isothermal-isobaric (NPT) ensemble at 25°C and 1 atm using a Langevin thermostat and Monte Carlo barostat (70–72). The g-BAOAB integrator was used with an integration timestep of 2 fs and trajectory frames were saved every 1000 steps (70,73). Bond lengths between hydrogens and heavy atoms were constrained using the SHAKE algorithm (74–77). Periodic boundary conditions were employed in conjunction with Particle Mesh Ewald (PME) electrostatics (78–80), to compute long-range electrostatic forces, and force-switched van der Waals (vdW) interactions (81). Nonbonded cutoffs were set to 10 Å, with force switching taking effect starting at 9 Å.

All explicit solvent calculations were conducted using the TIP3P water model (82). The CHARMM36 protein force field was used to represent the inosine and m^6^A Fabs, and the CHARMM36 nucleic acid force field was used to represent the RNA oligos (83–87). Modified ribonucleobase parameters were used to model noncanonical bases in the ribonucleoside (54). For the alchemical perturbations performed with λ-dynamics, ribonucleoside base mutations were represented using a hybrid multiple-topology approach (88). In the case of purine-to-purine mutations, analogous atoms in the shared core were harmonically restrained to one another using the Scaling of Constrained Atoms (SCAT) interface described previously (89).

### λ-Dynamics calculations

From 112 parameterized modified ribonucleobases available (54), a library of 48 bases, comprising 44 modified and 4 unmodified base candidates, were selected for *in silico* screening with λ-dynamics. Those with charged functional groups, bulky side chains, or modifications to the ribose sugar were excluded. Simulations were conducted for each of the 48 ribonucleosides with λ-dynamics to alchemically transform wild-type nucleoside bases (inosine or m^6^A) into a corresponding mutant base and compute relative differences in binding affinities. Prior to initiating λ-dynamics production sampling, appropriate biasing potentials must first be identified. The Adaptive Landscape Flattening (ALF) (49,90) algorithm was used to identify optimal biasing potentials to facilitate dynamic and frequent alchemical transitions between the perturbed bases. For each perturbation, ALF identified initial biases by first conducting one hundred simulations of 100 ps MD sampling, followed by 13 simulations of 1 ns each. These biases were then further refined via five replicate simulations of 5 ns each. With optimal biases identified, five independent production simulations of 25 ns were conducted, with an initial 5 ns of sampling removed from free energy determinations for equilibration. Ribonucleosides that unbound from the Fab binding site during λ-dynamics production sampling were labeled as unfavorable and were not pursued further. In all other cases, final ΔΔ*G*_bind_ values were calculated by Boltzmann reweighting the end-state populations to the original biases with WHAM (49,91). Uncertainties (*σ*) were calculated by bootstrapping the standard deviation of the mean across each of the five independent trials. From these results, modified oligonucleotides were selected for synthesis based on commercial availability.

### RNA oligonucleotide preparation

RNA oligonucleotides used for binding affinity measurements and crystallographic studies were synthesized on an ABI 394 DNA/RNA synthesizer (Applied Biosystems (ABI); Waltham, MA). m^6^A (10-3005-90; Glen Research; Sterline, VA), m^6^_2_A (ANP-8626; Chemgene; Wilmington, MA), and inosine (ANP-5680; Chemgene) modified RNA phosphoramidites; Biotin phosphoramidite (CLP-1517; Chemgene); and canonical RNA (A, ANP-5671; U, ANP-5674; C, ANP-6676; Chemgene) phosphoramidites were purchased from commercial sources. The canonical and modified phosphoramidites were concentrated to 0.1 M in acetonitrile. Coupling was carried out using a 5-benzylthio-1H-tetrazole (5-BTT) solution (0.25 M) as the catalyst. The coupling time was 650 seconds. 3% trichloroacetic acid in methylene chloride was used for the detritylation. Syntheses were performed on control pore glass (CPG-1000) immobilized with the appropriate nucleosides. All L-oligonucleotides were prepared with DMTr-on and in-house deprotected using AMA (1:1 v/v aqueous mixture of 30% w/v ammonium hydroxide and 40% w/v methylamine) for 15 minutes at 65°C. The RNA strands were additionally desilylated with Et_3_N•3HF solution to remove TBDMS groups. The 5’-DMTr deprotection was carried out using the commercial Glen-Pak purification cartridge (Glen Research). Purification was initially performed by the commercial Glen-Pak purification cartridge, followed by further purification with a 15% denaturing PAGE gel. The oligonucleotides were collected, lyophilized, desalted, re-dissolved in water, and then concentrated as appropriate for downstream experiments. Concentrations of the aqueous RNA samples were determined by their UV absorption at 260 nm, using the Thermo Scientific Nanodrop One Spectrophotometer. The theoretical molar extinction coefficients of these samples at 260 nm were provided by Integrated DNA Technologies.

### ELISA

Biotin-labeled, RNA oligos were diluted to 100 nM in ELISA blocking buffer (PBS, 0.05% Tween-20, 0.2 mg/ml bovine serum albumin (BSA, BP9706100; Fisher Scientific; Hampton, NH)), and 100 ul were incubated in clear, 96-well NeutrAvidin™ Coated Plates (PI15217; Pierce; Waltham, MA) overnight at 4^°^C. Two technical replicates were set for each RNA oligo. ELISA blocking buffer without oligo condition was used as a negative control. The plates were washed with PBS-T (PBS with 0.05% Tween-20) 3 times, and varying concentrations of recombinant mAb incubated in each well for 1 hour at room temperature (approximately 20^°^C). A no-mAb condition was used as a no primary antibody control. Plates were washed 3 times again with PBS-T and incubated with goat anti-mouse IgG conjugated to horseradish peroxidase (HRP, NBP2-30347H; Novus Biologicals; Centennial, CO) at 0.05 µg/ml in ELISA blocking buffer for 1 hour at room temperature (approximately 20^°^C). The plates were washed again with PBS-T and incubated with 50 ul of room temperature 1-Step Ultra TMB-ELISA Substrate Solution (PI34028; Pierce). After 15 minutes, the reaction was stopped with 50 ul of 2M Sulfuric Acid (A300S-500, Fisher Scientific). The plates were analyzed by 450 nm absorbance with a Synergy H1 microplate reader (BioTek Instruments; Winooski, VT). All ELISA experiments were replicated at least 3 times. The 3 cleanest runs were reported. Averages, standard deviations, and graphs were performed and made in GraphPad Prism version 10.1.1 for MacOS (GraphPad Software, Boston, MA).

## Acknowledgements

The authors thank Dr. Millie Georgiadis for help with crystal data collection and model building, and members of the Aoki, Vilseck, and Zhang Labs for their helpful discussion. Data collection and processing was also facilitated by Craig A. Bingman of the Department of Biochemistry Collaborative Crystallography Core and the University of Wisconsin, which is supported by user fees and the department. X-ray diffraction data were collected on Advanced Photon Source beamline LRL-CAT 31-ID and the European Synchrotron Radiation Facility (ESRF) ID30B operated by the Life Sciences Collaborative Access Team (LS-CAT). This research used resources of the Advanced Photon Source, a US Department of Energy (DOE) Office of Science User Facility operated for the DOE Office of Science by Argonne National Laboratory under Contract No. DE-AC02-06CH11357. Use of the Lilly Research Laboratories Collaborative Access Team (LRL-CAT) beamline at Sector 31 of the Advanced Photon Source was provided by Eli Lilly Company, which operates the facility. The authors acknowledge the Indiana University Pervasive Technology Institute for providing supercomputing and storage resources that have contributed to the research results reported within this paper. S.T.A, W.Z., and J.Z.V. received start-up funds from the Indiana University School of Medicine and its Precision Health Initiative (PHI). J.Z.V. gratefully acknowledges the National Institutes of Health (NIH) for financial support through grant R35GM146888. S.T.A. is funded by the NIH/NIGMS (R35GM142691) and an Indiana University Research Support Funds Grant (RSFG).

## Supplemental Figure Captions

**Table S1.** Data collection and refinement statistics for the inosine and m^6^A antibody crystal structures.

|  | **Inosine Fab w/**<br>**Inosine (PDB ID:**<br>**8SIP)** | **m6A Fab only**<br>**(PDB ID: 8TCA)** | **m6A Fab w**<br>**ligand (PDB ID:**<br>**8VEV)** |
| --- | --- | --- | --- |
| **Wavelength** | 0.9793 | 0.9793 | 0.8731 |
| **Resolution**<br>**range** | 57.04 - 1.94<br>(2.009 - 1.94) | 70 - 2.02 (2.092<br>- 2.02) | 48.23 - 3.06 (3.18<br>- 3.06) |
| **Space group** | P 1 | P 43 21 2 | P 21 21 21 |
| **Unit cell** | 39.8174 49.0903<br>57.3853 83.8419<br>88.8169 89.6813 | 79.906 79.906<br>145.127 90 90 90 | 83.64 128.377<br>150.476 90 90 90 |
| **Total reflections** | 218254 (14586) | 230543 (22581) | 203373 (23011) |
| **Unique**<br>**reflections** | 30888 (2931) | 31198 (2999) | 31244 (3399) |
| **Multiplicity** | 7.1 (4.9) | 7.4 (7.3) | 6.5 (6.8) |
| **Completeness**<br>**(%)** | 96.20 (91.25) | 98.24 (96.71) | 99.90 (100.00) |
| **Mean I/sigma(I)** | 6.33 (3.17) | 9.71 (1.08) | 16.80 (4.00) |
| **Wilson B-factor** | 26.61 | 27.17 | 78.19 |
| **R-merge** | 0.1901 (1.506) | 0.2183 (2.290) | 0.078 (0.416) |
| **R-meas** | 0.2027 (1.642) | 0.2341 (2.466) | 0.085 (0.451) |
| **R-pim** | 0.06938 (0.6427) | 0.08188 (0.8885) | 0.0330 (0.1720) |
| **CC1/2** | 0.991 (0.586) | 0.996 (0.546) | 0.998 (0.963) |
| **CC\*** | 0.998 (0.86) | 0.999 (0.84) | 1.000 (0.990) |
| **Reflections used**<br>**in refinement** | 30804 (2931) | 31042 (2999) | 31207 (3391) |
| **Reflections used**<br>**for R-free** | 1554 (154) | 1543 (162) | 1307 (142) |
| **R-work** | 0.2043 (0.2460) | 0.1807 (0.3043) | 0.2238 (0.3156) |
| **R-free** | 0.2454 (0.2914) | 0.2238 (0.3201) | 0.2531 (0.3100) |
| **CC(work)** | 0.947 (0.722) | 0.968 (0.777) | 0.928 (0.846) |
| **CC(free)** | 0.929 (0.494) | 0.952 (0.735) | 0.896 (0.865) |
| **Number of non-hydrogen atoms** | 3531 | 3562 | 9807 |
| **macromolecules** | 3277 | 3239 | 9649 |
| **ligands** | 19 | 36 | 108 |
| **solvent** | 235 | 287 | 50 |
| **Protein residues** | 426 | 425 | 1267 |
| **RMS(bonds)** | 0.008 | 0.009 | 0.002 |
| **RMS(angles)** | 1.10 | 1.18 | 0.56 |
| **Ramachandran favored (%)** | 97.62 | 97.61 | 95.34 |
| **Ramachandran allowed (%)** | 2.38 | 2.39 | 4.58 |
| **Ramachandran outliers (%)** | 0.00 | 0.00 | 0.08 |
| **Rotamer outliers (%)** | 0.80 | 0.54 | 0.09 |
| **Clashscore** | 6.46 | 1.70 | 2.09 |
| **Average B-factor** | 32.66 | 33.80 | 91.07 |
| **macromolecules** | 32.19 | 33.22 | 91.26 |
| **ligands** | 32.70 | 53.08 | 81.28 |
| **solvent** | 39.15 | 37.94 | 75.58 |
| **Number of TLS groups** | 1 | 9 | 6 |

**Table S2.** Complete table of λ-dynamics results for inosine antibody screening with the RNA library. RNA chemical structures available in **Fig S3**. Relative binding free energy, ΔΔ*G*_bind_. Standard deviation, ±*σ*. Unbound, u.b. Not specified due to bad sampling, n.s. Entries corresponding to favorable modifications (ΔΔ*G*_bind_ ≤ −0.7 kcal/mol) are emphasized in bold italics. Patch name from Xu et al., 2016.

| Modified Base | Patch Name | $\Delta\Delta G_{\text{bind}}$ | $\pm \sigma$ |
| --- | --- | --- | --- |
| A | ADE | -0.112 | 0.172 |
| $m^2A$ | 2MA | u.b. | u.b. |
| $m^6A$ | 6MA | 1.726 | 0.186 |
| $m^6_2A$ | M6A | 0.605 | 0.231 |
| $m^8A$ | 8MA | 2.721 | 0.389 |
| $m^1I$ | 1MI | 0.783 | 0.138 |
| I | INO | 0.089 | 0.024 |
| $ms^2m^6A$ | SMA | -0.037 | 0.335 |
| $ac^6A$ | 6AA | u.b. | u.b. |
| $i^6A$ | 6IA | n.s. | n.s. |
| $ms^2i^6A$ | MIA | 0.598 | 0.352 |
| $ms^2io^6A$ | SIA | u.b. | u.b. |
| $io^6A$ | HIA | 1.868 | 0.389 |
| G | GUA | -0.642 | 0.134 |
| $m^1G$ | 1MG | u.b. | u.b. |
| $m^2G$ | 2MG | 0.286 | 0.168 |
| $m^2_2G$ | M2G | 1.057 | 0.262 |
| **preQ0** | **DCG** | **-1.737** | **0.187** |
| *imG-14* | DWG | -0.005 | 0.319 |
| *imG* | IMG | u.b. | u.b. |
| **imG2** | **IWG** | **-1.164** | **0.331** |
| *mimG* | MWG | u.b. | u.b. |
| **U** | **URA** | **-0.926** | **0.145** |
| D | H2U | u.b. | u.b. |
| **mo<sup>5</sup>U** | **MOU** | **-1.656** | **0.199** |
| $m^5s^2U$ | 52U | u.b. | u.b. |
| $m^5D$ | MDU | 0.707 | 0.161 |
| **$\psi$** | **PSU** | **-1.985** | **0.490** |
| **$m^3\psi$** | **3MP** | **-1.174** | **0.446** |
| $m^3U$ | 3MU | -0.146 | 0.320 |
| **$s^4U$** | **4SU** | **-1.702** | **0.181** |
| **$m^5U$** | **5MU** | **-1.340** | **0.159** |
| **ho<sup>5</sup>U** | **5HU** | **-1.613** | **0.162** |
| $s^2U$ | 2SU | -0.644 | 0.233 |
| **$m^1\psi$** | **1MP** | **-0.896** | **0.112** |
| **cnm<sup>5</sup>U** | **CYU** | **-2.147** | **0.233** |
| **mcm<sup>5</sup>s<sup>2</sup>U** | **70U** | **-1.379** | **0.330** |
| *mchm<sup>5</sup>U* | CMU | -0.612 | 0.381 |
| **ncm<sup>5</sup>U** | **BCU** | **-1.683** | **0.393** |
| **mcm<sup>5</sup>U** | **OCU** | **-1.307** | **0.312** |
| *mcmo<sup>5</sup>U* | OEU | -0.491 | 0.396 |
| C | CYT | u.b. | u.b. |
| $m^5C$ | 5MC | u.b. | u.b. |
| $ac^4C$ | 4AC | u.b. | u.b. |
| $m^4C$ | 4MC | u.b. | u.b. |
| $f^5C$ | 5FC | 0.596 | 0.265 |
| $hm^5C$ | HMC | u.b. | u.b. |
| $s^2C$ | 2SC | u.b. | u.b. |

"Patch Name" = 3-letter name assigned by Xu et al. (2016)
Modifications with $\Delta\Delta G \leq -0.7$ kcal/mol in bold italics.

**Table S3.** Complete table of λ-dynamics results for m^6^A antibody screening with the RNA library. RNA chemical structures available in **Fig S3**. Relative binding free energy, ΔΔ*G*_bind_. Standard deviation, ±*σ*. Unbound, u.b. Not specified due to bad sampling, n.s. Entries corresponding to favorable modifications (ΔΔ*G*_bind_ ≤ −0.7 kcal/mol) are emphasized in bold italics. Patch name from Xu et al., 2016.

| Modified Base | Patch Name | $\Delta\Delta G_{\text{bind}}$ | $\pm \sigma$ |
| --- | --- | --- | --- |
| A | ADE | 0.300 | 0.165 |
| *m*<sup>2</sup>A | 2MA | 3.084 | 0.156 |
| *m*<sup>6</sup>A | 6MA | -0.034 | 0.032 |
| ***m*<sup>6</sup><sub>2</sub>A** | **M6A** | **-2.091** | **0.052** |
| *m*<sup>8</sup>A | 8MA | 4.432 | 0.337 |
| *m*<sup>1</sup>I | 1MI | n.s. | n.s. |
| I | INO | 6.247 | 0.301 |
| *ms*<sup>2</sup>*m*<sup>6</sup>A | SMA | 2.634 | 0.209 |
| ***ac*<sup>6</sup>A** | **6AA** | **-1.156** | **0.197** |
| *i*<sup>6</sup>A | 6IA | 1.624 | 0.491 |
| *ms*<sup>2</sup>*i*<sup>6</sup>A | MIA | 3.587 | 0.434 |
| *ms*<sup>2</sup>*io*<sup>6</sup>A | SIA | n.s. | n.s. |
| *io*<sup>6</sup>A | HIA | n.s. | n.s. |
| G | GUA | 4.881 | 0.471 |
| *m*<sup>1</sup>G | 1MG | 3.029 | 0.521 |
| *m*<sup>2</sup>G | 2MG | u.b. | u.b. |
| *m*<sup>2</sup><sub>2</sub>G | M2G | n.s. | n.s. |
| *preQ0* | DCG | 4.229 | 0.283 |
| *imG*-14 | DWG | u.b. | u.b. |
| *imG* | IMG | n.s. | n.s. |
| *imG*2 | IWG | u.b. | u.b. |
| ***mimG*** | **MWG** | **-2.437** | **0.441** |
| U | URA | u.b. | u.b. |
| D | H2U | 2.841 | 0.242 |
| *mo*<sup>5</sup>U | MOU | 1.861 | 0.459 |
| *m*<sup>5</sup>*s*<sup>2</sup>U | 52U | 3.817 | 0.504 |
| *m*<sup>5</sup>D | MDU | 1.520 | 0.25 |
| ψ | PSU | 3.564 | 0.531 |
| *m*<sup>3</sup>ψ | 3MP | n.s. | n.s. |
| *m*<sup>3</sup>U | 3MU | n.s. | n.s. |
| *s*<sup>4</sup>U | 4SU | n.s. | n.s. |
| *m*<sup>5</sup>U | 5MU | 1.270 | 0.519 |
| *ho*<sup>5</sup>U | 5HU | 0.965 | 0.305 |
| *s*<sup>2</sup>U | 2SU | 2.830 | 0.705 |
| *m*<sup>1</sup>ψ | 1MP | 3.044 | 0.663 |
| *cnm*<sup>5</sup>U | CYU | 2.396 | 0.282 |
| *mcm*<sup>5</sup>*s*<sup>2</sup>U | 70U | n.s. | n.s. |
| *mchm*<sup>5</sup>U | CMU | 2.686 | 0.235 |
| *ncm*<sup>5</sup>U | BCU | n.s. | n.s. |
| *mcm*<sup>5</sup>U | OCU | 1.113 | 0.402 |
| *mcmo*<sup>5</sup>U | OEU | 3.469 | 0.527 |
| C | CYT | 5.033 | 0.708 |
| *m*<sup>5</sup>C | 5MC | n.s. | n.s. |
| ***ac*<sup>4</sup>C** | **4AC** | **-1.199** | **0.325** |
| *m*<sup>4</sup>C | 4MC | 2.159 | 0.287 |
| *f*<sup>5</sup>C | 5FC | 2.275 | 0.422 |
| *hm*<sup>5</sup>C | HMC | u.b. | u.b. |
| *s*<sup>2</sup>C | 2SC | u.b. | u.b. |

"Patch Name" = 3-letter name assigned by Xu et al. (2016)
Modifications with $\Delta\Delta G \leq -0.7$ kcal/mol in bold italics.

**Fig S1.**
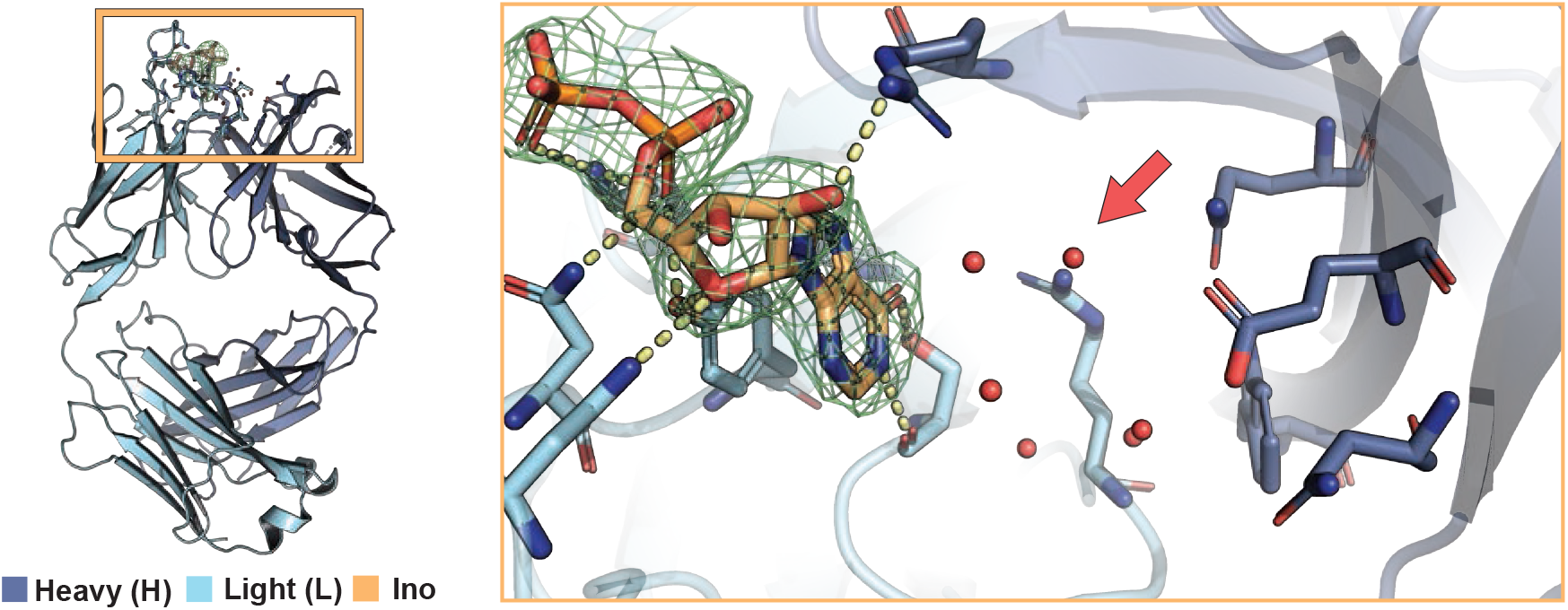
A previously published poly-inosine antibody has a large binding pocket that may accommodate multiple nucleobases. Overview (left) and magnified image (right) of the poly-inosine antibody fragment (PDB ID: 1MRD) binding pocket. An inosine mononucleotide (orange) was modeled into the missing ligand density (green). Heavy chain residues (H) in dark blue and light chain residues (L) in light blue. Water molecules substituting for the potential second mononucleotide are depicted as red spheres, indicating the potential space to bind a second nucleobase. Interacting amino acids include heavy chain residue Arg96 and light chain residues Asn28, Asn30, Tyr32, Lys50, and Ser91. The extended binding pocket (red arrow) includes light chain residue Arg96 and heavy chain residues Gln35, Trp47, Glu50, and Asn58.

**Fig S2.**
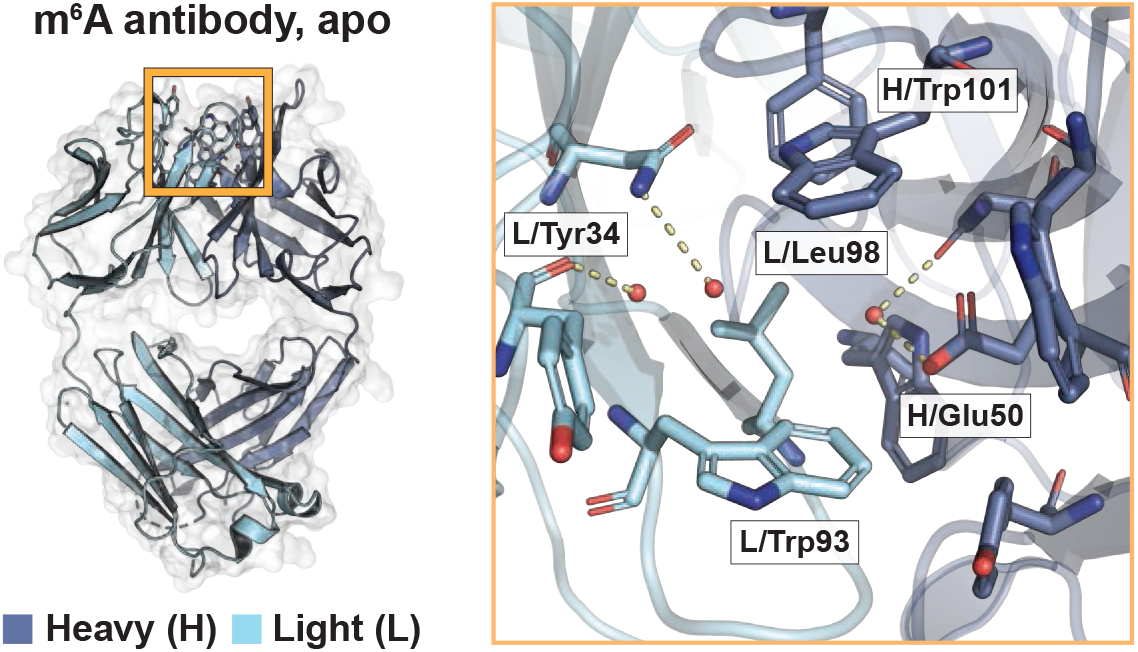
Crystal structure of the m^6^A Fab apo-form to 2.05 Å (PDB ID: 8TCA). Critical binding pocket amino acids discussed in the main text are labeled. Heavy chain residues (H) are represented in dark blue, light chain residues (L) in light blue, and waters as red spheres. Depicted binding pocket amino acids match those of the m^6^A Fab holo-form (**Fig 1A**).

**Fig S3.**
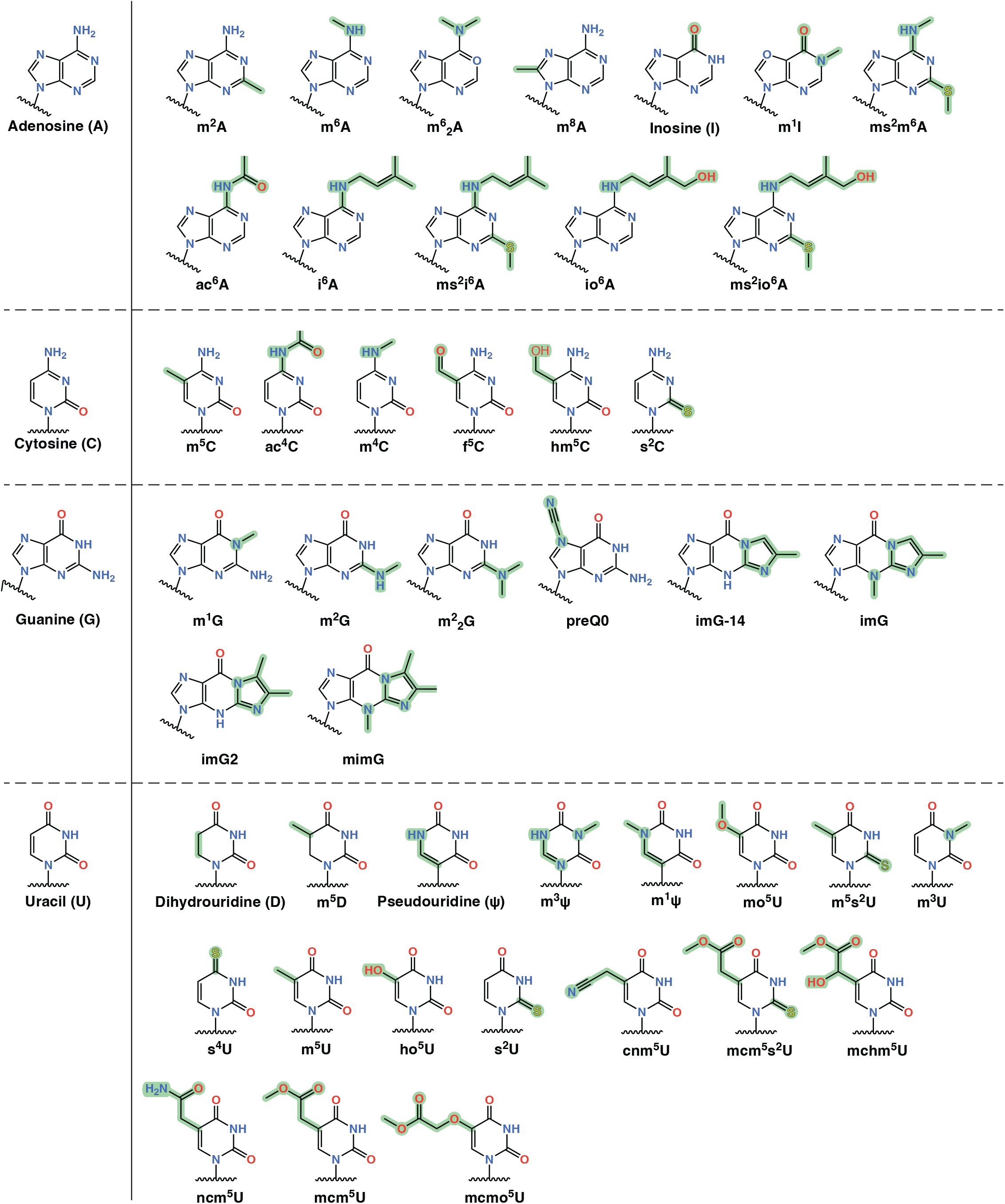
Chemical library of ribonucleoside bases. The library includes the 4 canonical ribonucleobases (A, C, G, and U) and 44 naturally occurring modified derivatives (12 As, 6 Cs, 8 Gs, and 18 Us). Differences between each modification and its respective canonical base are highlighted in green.

**Fig S4.** Molecular dynamics simulation movie example of the m^6^A antibody with a bound nucleoside target. The m^6^A Fab binds tightly to m^6^_2_A. Movie made in Pymol (Schrödinger, Inc.).

**Fig S5.** Molecular dynamics simulation movie example of the m^6^A antibody with an unbinding nucleoside target. The m^6^A Fab unbinds from uridine. Movie made in Pymol (Schrödinger, Inc.).

**Fig S6.**
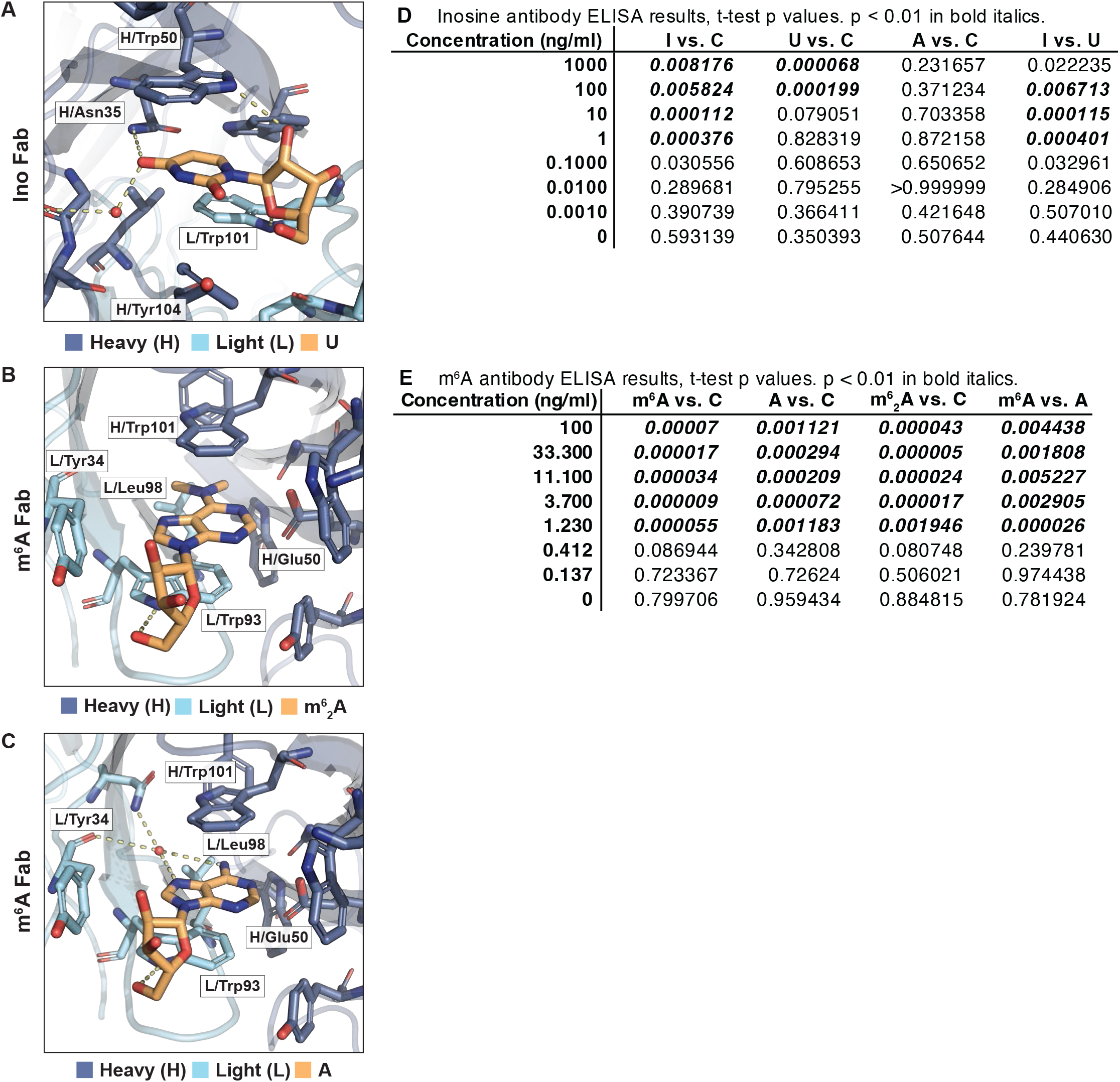
Structural models of inosine and m^6^A antibodies bound to representative off-target RNAs. (**A**) Magnified binding site of the inosine antibody fragment in complex with uridine. (**B-C**) Magnified binding site of the m^6^A antibody fragment in complex with (**B**) m^6^_2_A or (**C**) adenosine (A) Heavy chain residues (H) are represented in dark blue, light chain residues (L) in light blue, and the off-target nucleoside in orange. Critical amino acid contacts labeled. (**D-E**) Table of t-test p-value statistics for (**D**) inosine and (**E**) m^6^A antibody ELISA binding assay results reported in **Fig 5**. p-values < 0.01 in bold.

